# Male song sparrows modulate their aggressive signaling in response to plumage signals: experiments with 3-D printed models

**DOI:** 10.1101/753772

**Authors:** Michelle L. Beck, Ҫağlar Akҫay, Kendra B. Sewall

**Affiliations:** Department of Biological Sciences, Virginia Tech, Blacksburg, VA 24061, USA; Department of Biology, Rivier University, Nashua, NH 03060,USA; Department of Psychology, Koç University, Istanbul, 34450, Turkey

**Keywords:** 3-D printed model, Intrusion experiment, Male competition, Melanin, Plumage colouration, Song sparrow, Taxidermic mount, Territorial aggression

## Abstract

Competitive interactions among conspecifics are often resolved by assessing signals that honestly indicate individual fighting ability or dominance. In territorial species, signals of competitive ability are thought to function primarily during the early stages of territory establishment, but recent evidence suggests that these signals continue to influence interactions with floaters and neighbors well after territory establishment. Here, we examine the influence of the extent of chest spotting displayed by an intruding male on the response of territorial male song sparrows. We exposed males to 3-D printed models with large or small spotting area coupled with conspecific playback and recorded their behavior. We also assessed the response of a subset of males to both the 3-D printed models and a traditional, taxidermic mount to ensure the 3-D models were a realistic stimulus. We found no differences in the number of attacks or proximity to the model due to spotting area. However, territorial males produced more soft songs and tended to sing fewer loud songs, both of which predict attack in our population, in response to the model with less chest spotting. One possibility is that males with less chest spotting elicit a stronger response because they are seen as a greater threat. Based on our previous findings in this system, we think it is more likely that models with less chest spotting are perceived as subordinate and therefore easier to defeat, leading to a stronger response by territory holders. We found males were equally likely to attack 3-D printed models and a taxidermic mount but signaled more aggressively during trials with the taxidermic mount than the 3-D printed models. This suggests that birds recognized the 3-D models as meaningful stimuli but that the use of 3-D printed models should be validated through comparison to a traditional taxidermic mount when possible.

---

Intrasexual competition for access to mates or resources is a powerful selective force in many animals. Competitive interactions can be costly, which can lead to the evolution of signals that indicate dominance or fighting ability that can resolve these interactions without physical fights (Smith & Parker, 1976; Smith & Price, 1973). For signaling systems to be stable, the signaler must benefit from the receivers’ response and the receivers must benefit from responding to the information conveyed by the signal (Searcy & Nowicki, 2005; Smith & Parker, 1976). Additionally the signal must be honest, meaning that cheating - exaggerating one’s level of fighting ability - should occur infrequently in the population (Webster, Ligon, & Leighton, 2018). Reliable signals of dominance or resource holding potential are thought to be most likely to evolve in species that live in groups or in species in which frequent challenges occur among unfamiliar conspecifics (Rohwer, 1975, 1982; Senar, 2006). In this context, the honesty of the signal is maintained by social costs because individuals signaling above their rank are challenged and defeated repeatedly by group members. Once individuals are familiar with each other, prior experience is expected to influence the outcome of competitive interactions to a greater extent than a signal of fighting ability (Chaine, Shizuka, Block, Zhang, & Lyon, 2018; Lemel & Wallin, 1993; Senar, 2006; Vedder, Schut, Magrath, & Komdeur, 2010).

In territorial species, signals of competitive ability are thought to only function during the initial stages of territory establishment when individuals are unfamiliar with each other (Lemel & Wallin, 1993; Part & Qvarnstrom, 1997; Senar, 2006). Ornaments, such as bright coloration, are traits that act as signals of mate quality or fighting ability, but are not used in combat with other males. For instance, natural or experimentally induced variation in male ornaments is related to their ability to acquire nest sites (Part & Qvarnstrom, 1997; Pryke & Andersson, 2003a; Siefferman & Hill, 2005) or secure a high quality territory (Keyser & Hill, 2000). However, following territory establishment, most social interactions will occur between neighbors with whom individuals are familiar and possession of a territory confers an ownership advantage thought to render a phenotypic signal of resource holding potential irrelevant (Rohwer, 1982; Senar, 1999). Despite this, recent studies indicate that male ornaments continue to function post-territory establishment as more ornamented territory holders often experience fewer intrusions from floaters and neighbors during the breeding season (Chaine & Lyon, 2008; Cline, Hatt, Conroy, & Cooper, 2016; Pryke & Andersson, 2003a; Pryke, Lawes, & Andersson, 2001). Additionally, male territory holders modulate their response to conspecific intruders based on the intruder’s ornamentation and may either respond less strongly to more ornamented males or may respond more strongly to more ornamented individuals or to individuals that have ornamentation similar to their own (Chaine & Lyon, 2008; Martin et al., 2016; Pryke et al., 2001). These findings suggest that ornaments remain important signals of fighting ability or resource holding potential throughout the breeding season. Nevertheless, relatively few studies have examined the utility of ornaments in mediating social interactions post-territory establishment in species or populations with few floaters (but see Cline et al. 2016).

Conspicuous colouration is a signal used to mediate aggressive interactions in a variety of taxa including insects (Tibbetts & Dale, 2004), reptiles (Ligon & McGraw, 2016; Mafli, Wakamatsu, & Roulin, 2011; Martin et al., 2016; Seddon & Hews, 2016), fish (Johnson & Fuller, 2015; Schweitzer, Motreuil, & Dechaume-Moncharmont, 2015), and has been especially well studied in birds (reviewed in Senar, 2006; Tibbetts & Safran, 2009). In birds, conspicuous colouration can be produced by feather microstructure or by the deposition of pigments, such as carotenoids or melanins in feathers. Melanin-based colouration produces brown, black, and reddish plumage and the role of melanin-based traits in mediating competitive interactions has frequently been assessed. A number of studies have found larger or darker melanin-based plumage patches are associated with higher social status in flocks (Rohwer, 1975, 1977) and greater fighting ability or dominance (Chaine & Lyon, 2008; Chaine, Tjernell, Shizuka, & Lyon, 2011; Dunn, Whittingham, Freeman-Gallant, & DeCoste, 2008; Gonzalez, Sorci, Smith, & de Lope, 2002; Santos, Scheck, & Nakagawa, 2011; Tarof, Dunn, & Whittingham, 2005). In territory holding species, darker males respond more strongly to model intruders and darker model intruders are subject to more attacks and are approached more quickly than lighter models (Chaine & Lyon, 2008). But, darker territory holding males are themselves subject to more intrusions than lighter males (Chaine & Lyon, 2008). However, the consistency of some of these relationship has recently been questioned and much of this research has focused on the house sparrow (Passer domesticus, Kingma et al., 2008; Sanchez-Tojar et al., 2018). Further research on melanin-based ornaments in a greater variety of species and in species that are territorial is needed.

One approach used to determine if ornaments mediate aggressive interactions is to present conspecifics with one or several taxidermic mounts that vary in the size or reflectance of a colour patch and record the response of the focal individual (Chaine & Lyon, 2008; Coady & Dawson, 2013; Korsten, Dijkstra, & Komdeur, 2007; Pryke et al., 2001). Taxidermic mounts are advantageous because they provide a consistent stimulus which can permit focusing solely on the effects of colouration without confounding changes in behavior as can be seen when free living individuals are manipulated or live decoys are used (Scriba & Goymann, 2008). However, the use of taxidermic mounts can be problematic. Taxidermic mounts necessitate collecting multiple individuals or attempting to find individuals that died from natural causes (Chaine & Lyon, 2008; Laubach, Blumstein, Romero, Sampson, & Foufopoulos, 2013). Further, mounts are likely to be subjected to attack during trials leading to cumulative damage over experiments or they must be protected in some way which leads to a less natural stimulus. Thus, a method of producing accurate models that are relatively easy to manipulate the ornamentation of or replace when needed would be ideal for studies focused on colouration in a variety of taxa. Recent advances in 3-D printing have allowed biologists to quickly and cheaply produce models for use in field research (Bentz, Philippi, & Rosvall, 2019; Fan et al., 2018; Igic et al., 2015). The advantages of 3-D printing over taxidermic models is that many copies, standardized in size and shape that are resistant to attacks can be produced. While several studies have utilized 3-D printed models in behavioral assays, few have compared the response of the same birds to both 3-D printed and traditional taxidermic models to determine if males respond similarly to both stimuli (but see Bentz et al., 2019 for a comparison of responses to 3-D models and live decoys with data gathered on different individuals).

Male song sparrows (*Melospiza melodia*) are territorial and possess brown spotting on their breast that ranges from reddish-brown to dark brown and varies in area (hereafter spotting area). The spotting is prominently displayed on the chest and is similar to spotting that acts as a signal in other species (Grunst & Grunst, 2015), but relatively little research has focused on the function of chest spotting in song sparrows. Male song sparrows occur in both urban and rural habitats and males in urban habitats display more extensive spotting and greater territorial aggression than males in rural habitat (Beck, Davies, & Sewall, 2018; Davies & Sewall, 2016; Foltz et al., 2015). In rural (but not urban) habitats, territorial males with less extensive spotting area are more aggressive during a simulated territorial intrusion than males with more extensive spotting (Beck et al., 2018). This finding is interesting given that in other species, birds with larger melanin-based ornaments are generally found to display greater territorial aggression (reviewed in Santos et al., 2011; Senar, 2006).

In this study, our goal was to determine how chest spotting influences aggressive interactions between male song sparrows. To do this, we presented territorial males with 3-D printed model song sparrows painted with large or small spotting area on their chests, while standardizing for spotting reflectance. A second aim of our study was to verify that 3-D printed models can be used to assess territorial aggression. To this end, we also presented a subset of males with a taxidermic mount of a song sparrow in addition to the 3-D models to compare responses to models and mounts.

## Methods

### Subjects and study sites

We studied song sparrows in rural and urban habitats located in Montgomery County, VA from 15-18 May 2017. For the present study the subjects were 14 male song sparrows living on the Virginia Tech campus (urban habitat) and 14 males living in Heritage Park and Stroubles Creek Stream Restoration Site (rural habitats). The details of the sites, including levels of urbanization can be found in Davies et al. (2018). Seven of the urban birds and 1 of the rural birds were banded, the rest of them were non-banded. The trials were conducted on consecutive days approximately 24 hours apart to ensure the same male was sampled each time. Each rural subject was tested twice in a counterbalanced order: once with a small spotting area model and once with a model with large spotting area. Eleven of the urban subjects were tested three times, once with a taxidermic mount, once with a small spotting area model and once with a large spotting area model. The remaining three were tested two times because they did not appear for the third trial: two of them were tested with the large and small-spotting area model and one with the mount and the small spotting area model. One more male in the urban habitat was tested only once, disappearing before the second trial. We included all males that had at least two trials in our comparisons. We only tested urban males with a mount, because we expected that if there is a difference in response to the 3-D models relative to the taxidermic mount it would be in the direction of a lower response and using the more aggressive urban males gave us a better chance to detect that difference.

### 3-D printing models

The original file for the model was made in Autodesk 123D app from a series of pictures of a plastic bird model, and then edited using Autodesk Meshmixer and saved as a .stl file. The model was designed such that there were no legs but it could be placed on the belly to stand upright (see Fig. 1 and the supplementary .stl file). We printed 6 models and then painted the models using acrylic paint to imitate the song sparrow plumage. Three of the models were painted with small spotting area (mean ± SE, 119.28 ± 0.880 mm^2^, range 117.5-120.3 mm^2^) and the other three were painted with a large spotting area 283.9 ± 45.61 mm^2^, range 238.3-329.5 mm^2^). We also used a taxidermic mount of a song sparrow to compare responses to the model (badge area 259.62 mm^2^).

**Figure 1.**
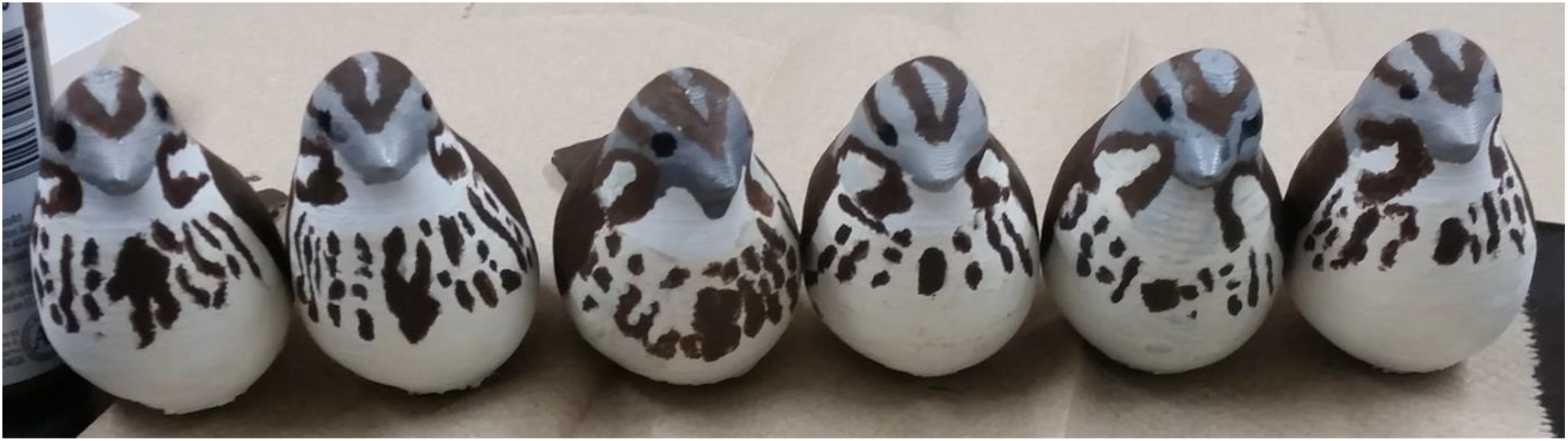
The six 3-D printed model song sparrows used in the behavioral trials. The three models on the left have large spotting area and the three on the right small spotting area.

#### Stimuli and trial procedure

The stimulus songs were recorded from song sparrows in Blacksburg, VA or Radford, VA using a Sennheiser directional microphone (ME66/K6) and Marantz PMD 660 or 661 solid state recorder. We selected stimulus songs based on the quality of recording. We added a silent period at the end of the song to create a 10 second playback clip using Syrinx (John Burt, Seattle, WA). We made 25 different stimuli tapes from 13 different males. Each subject received a single stimulus song type for all experimental conditions. The stimuli used for each subject came from birds that lived at least 2 km away from the subject.

The trials started when a singing male was located and a brief period of playback was used to identify the center of the male’s territory. The experimenter placed a tripod in the center of a territory near natural perches and placed a wireless speaker (VictSing model C6) on the tripod at a height of about 1 m. The taxidermic mount or the 3-D printed model was placed on top of the tripod above the speaker and covered with a cloth. The speaker was connected to a smartphone via Bluetooth, and the experimenter controlled the playback at a distance of about 15-20m.

With the model or mount covered, the behavior of the male was recorded for three minutes after the first response to the playback to obtain a baseline aggressive response. Following the pre-model period, we paused the playback and removed the cloth by walking over to the tripod and then restarted the playback. This model period of the trial lasted from the first time the subject entered within a 5m radius of the model/mount (as we wanted to ensure that the subjects saw the model or the mount) until either an attack (physically touching the model or mount) or 5 minutes has elapsed.

#### Response measures

We recorded the trial using the same recording equipment as above, narrating the behavior of the subject. We noted two aggressive behaviors, attacks and distance to the speaker, and three aggressive signaling behaviors, loud songs, soft songs (low amplitude songs), and wing waves. Loud song and soft songs were classified in the field by either CA or MLB; this method has been shown to be reliable in this species (Anderson, Searcy, Peters, & Nowicki, 2008). Soft songs and wing waves have been shown to be reliable signals of aggression (i.e. predicting a subsequent attack on a taxidermic mount) in multiple populations of this species, including the present one (Açkay, Beck, & Sewall, in review Akçay, Tom, Campbell, & Beecher, 2013; Searcy, Akçay, Nowicki, & Beecher, 2014; Searcy, Anderson, & Nowicki, 2006).

We scanned and annotated the trial recordings using Syrinx to extract the following information: Proportion of the trial spent within 1m of the speaker and counts of loud songs, soft songs and wing waves for each period. We converted the counts into rates by dividing the counts by the duration of the period to account for unequal observation durations.

#### Data analysis

The response variables were not normally distributed and we used non-parametric tests throughout. We first asked whether the models elicited different responses than a taxidermic mount in urban birds. For this, we compared the proportion of the trial within 1 m, loud song rates, soft song rates and wing waves in the urban subjects that received the mount treatment as well as at least one 3-D printed model (n=12). Eleven of the 12 subjects received both large and small spotting area 3-D models. For these subjects we averaged their responses to these models and compared the responses to mount with the responses to the 3-D printed models with a Wilcoxon signed-rank test. For individuals that received 3 trials, we used a Friedman’s test to determine if responses differed due to trial order.

Then we compared the responses to the small and large spotting area models using all the subjects that received both stimuli (n=27). We used a permutation test to test the main effect of condition (a within subject variable) and habitat (a between subject variable) and their interaction using the ezPerm function in the R package ez (Lawrence, 2016). Because behavioural studies frequently have issues with low statistical power, we did not perform a Bonferroni correction (Nakagawa, 2004).

#### Ethical note

This research adheres to the ABAS/ABS Guidelines for the Use of Animals in Research. All of our methods were approved by the Virginia Tech IACUC committee (BIOL 15-185). VA-DGIF (permit 48639), USGS bird banding lab (permit 23818) and the US Fish and Wildlife Service (MB08005B-0). We sampled 28, after-hatch year, wild, male song sparrows (*Melospiza melodia*). Many of these birds are found in urban areas where they commonly experience human disturbance. During observations, we remained 15-20 m away from the focal male which should have limited our effect on his behaviour and the trials were brief, only 8 minutes or until the male attacked the model. We did not capture males for this study (some were previously banded for studies in past years) and thus this was a minimally invasive project. Males that were previously captured were banded with one USGS metal band (size 1B) and 3 coloured leg bands (diameter 2.8 mm). These bands were not removed so that birds could be identified in future years. Leg bands are small, lightweight, and commonly used by ornithologists around the globe and should have minimal effect on a bird.

## Results

### Spotting area and male territorial aggression

The proportion of the trial spent within 1m of the 3-D models did not differ between the small- and large-spotting area models. However, habitat had a strong effect with urban birds spending more time within 1m of the models than rural birds as has been found in previous studies in this population (Davies & Sewall, 2016; Foltz et al., 2015). The interaction between habitat and condition was not significant (Table1, Fig 2a).

**Table 1.**
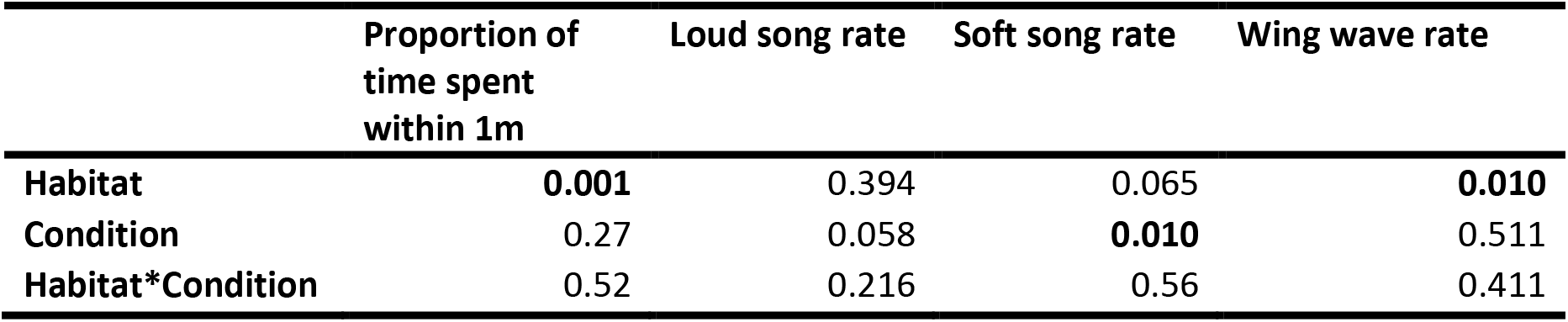
P-values from the permutation test on territorial aggression of male song sparrows during the presentation of 3-D printed models with large or small chest spotting in urban and rural habitats (1000 permutations).

|  | Proportion of<br>time spent<br>within 1m | Loud song rate | Soft song rate | Wing wave rate |
| --- | --- | --- | --- | --- |
| **Habitat** | **0.001** | 0.394 | 0.065 | **0.010** |
| **Condition** | 0.27 | 0.058 | **0.010** | 0.511 |
| **Habitat\*Condition** | 0.52 | 0.216 | 0.56 | 0.411 |

**Figure 2:**
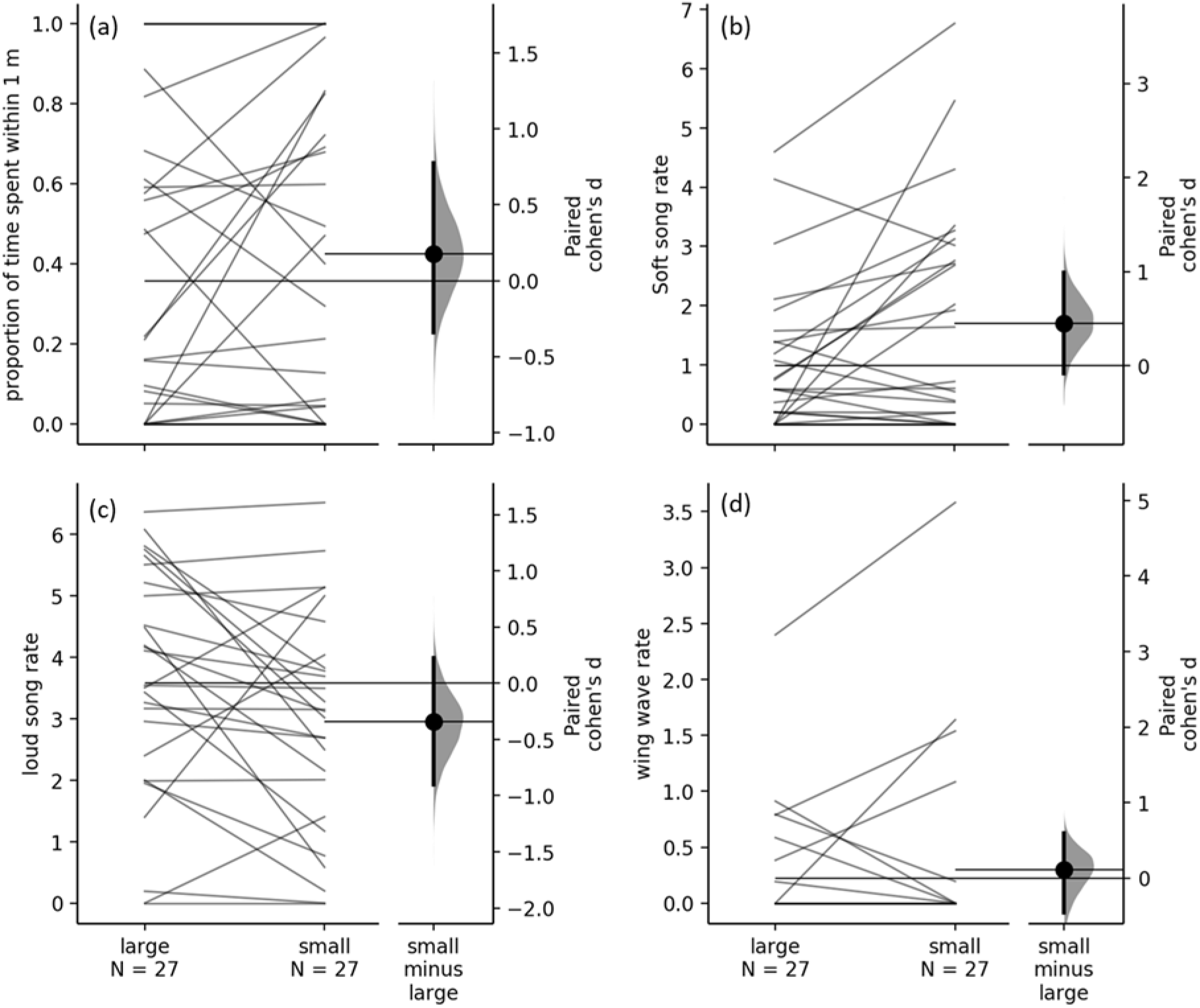
The responses of territorial male song sparrows to 3-D printed models with small or large spotting area in a) proportion of time spent within 1m, b) soft song rates, c) loud song rates, and d) wing wave rates. Rates are per minute. The lines are individual subjects.

In the signaling variables, there was a significant difference in rates of soft songs given in response to small and large spotting area models: subjects tended to give more soft songs in response to the models with small spotting area. The effect of habitat approached significance with urban birds tending to sing more soft songs, and the interaction effect was not significant (Table 1, Fig. 2b). Loud song rates showed a tendency to differ between conditions as well with subjects singing fewer loud songs to models with small spotting area (Fig. 2c). Finally, subjects did not differ in their wing wave rates between conditions, but urban birds gave significantly more wing waves (Fig. 2d, only one rural bird gave any wing waves).

Four subjects out of 27 (14.8%) attacked the model with small spotting area, whereas one subject (3.7%) attacked the large spotting area models. The difference was not significant by a chi-square test; χ^2^=1.98, p= 0.16. Two out of 11 subjects attacked the taxidermic mount (18.2%).

#### Response to 3-D printed models and the taxidermic mount

During the pre-model period, there were significant differences between the model and mount in proportion of time spent within 1 m (V=57,n=12,p=0.037): subjects spent significantly less time near the speaker in the mount trials than in the model trials before the model or mount was revealed. No other significant differences were detected for the pre-model period (all p ≥ 0.38). During the model presentation, there were significant differences between the responses to the taxidermic mount and the 3-D models. Subjects spent more time within 1m of the mount than the model (V=1, n=12, p = 0.01, Fig. 3a); sang more soft songs (V=13, n=12, p = 0.04, Fig. 3b), and more loud songs (V=67, n=12, p = 0.03, Fig. 3c) to the mount than to the 3-D models. Rates of wing waves did not differ significantly between the mount and the 3-D models (V=5, n=12, p = 0.15, Fig. 3d). The birds showed no signs of habituation as none of the response variables differed by trial number (all p > 0.15).

**Figure 3.**
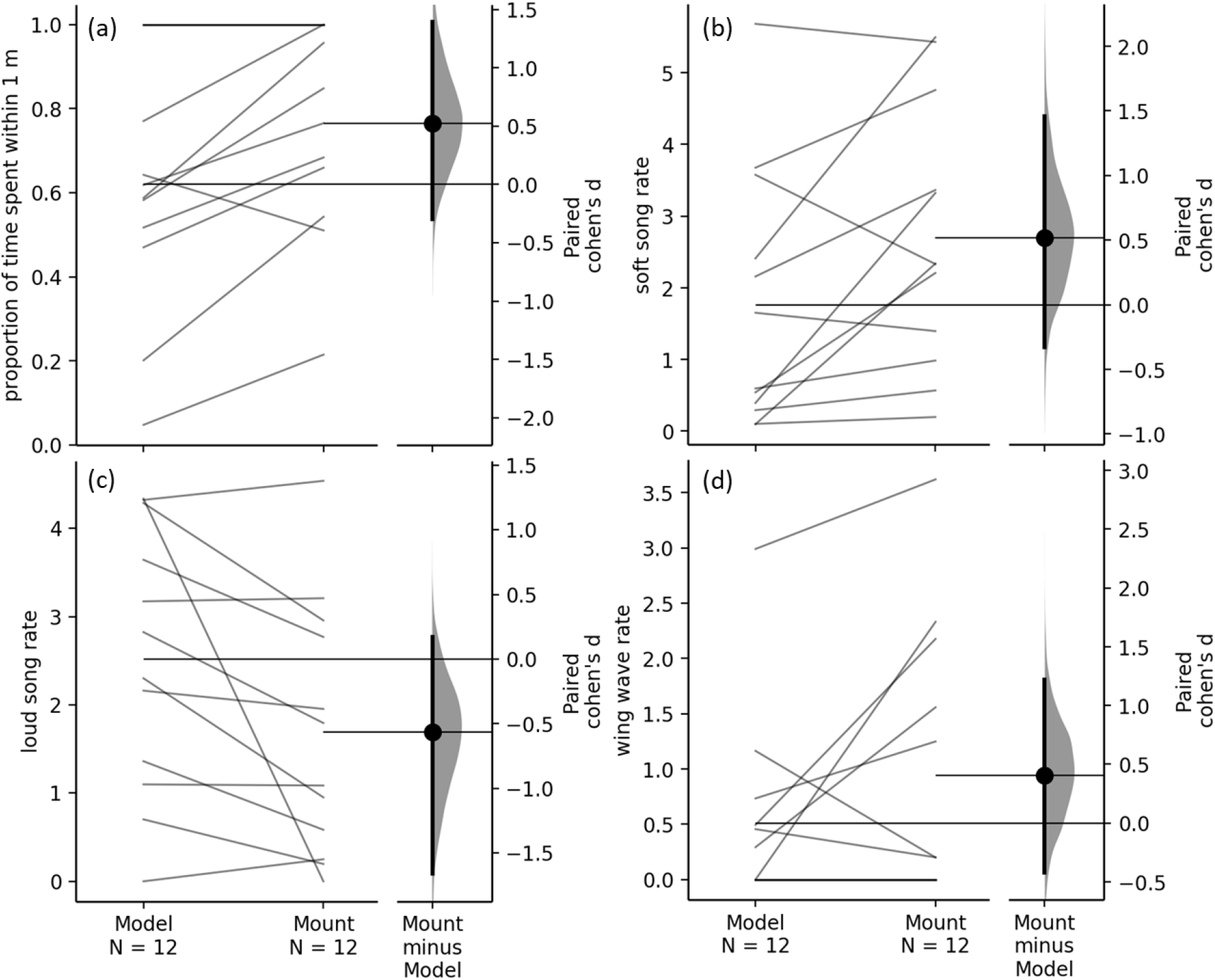
The responses of territorial male song sparrows to 3-D printed models and a taxidermic mount in a) proportion of time spent within 1m, b) soft song rates, c) loud song rates, and d) wing wave rates. Rates are per minute. The lines are individual subjects.

## Discussion

In this study we had two aims: 1) to determine whether male song sparrows respond differently to intruders based on the extent of chest spotting and 2) to determine whether a 3-D printed model can be effectively used to replace taxidermic mounts to study plumage signals. We found that when birds were presented with 3-D printed models with different sized spotting areas, they responded with more aggressive signaling towards the models with less chest spotting. We found that responses to the mount and 3-D printed models did differ significantly with the taxidermic mount eliciting a stronger aggressive response though the mount and 3-D models were attacked at similar rates.

### Spotting area as a signal of aggression

We expected to find a difference in aggressive and signaling behaviours in response to variation in spotting area. However, we only found a difference in soft songs and a trend for loud songs, but no differences in attack or proximity to the model. Because spotting area is a visual stimulus, relatively close approach may be necessary for assessment, leading to a lack of difference between treatments. Furthermore, the lack of behavioural response by the model may lead males to remain in close proximity to the model, irrespective of differences in spotting area. More soft songs were produced in response to the models with less chest spotting and soft songs are the most reliable signal of aggression in this species (Akçay et al., 2013; Searcy et al., 2006). Similarly, models with small spotting area tended to elicit lower rates of loud songs than models with large spotting area, and low rates of loud singing are predictive of physical attack in our population (Akçay et al. in review). Thus, these two findings can be interpreted as a difference in aggressive signaling, even though there was no difference in approach or attacks.

A difference in responses to chest spotting size is consistent with chest spotting serving as a signal reflecting status or resource holding potential in song sparrows. However, whether higher aggression towards males with small spotting area means that these are perceived to be a greater threat or are viewed as easier to defeat requires further testing. Subjects may respond with greater intensity to small spotting models because these represents a greater threat. Alternatively, subjects may respond more strongly to small spotting area models because these represent a lower threat which makes investment in aggressive behaviours less costly (in terms of risk of retaliation and injury) than it would be in response to a higher threat opponent with a large spotting area (Pryke & Andersson, 2003a; Searcy & Beecher, 2009). We think this latter explanation is more likely because urban male song sparrows have more extensive chest spotting and display greater territorial aggression than rural males (Beck et al., 2018; Davies & Sewall, 2016). The negative association between spotting area and territorial aggression we previously found in rural males may be the result of rural males with less chest spotting being frequently challenged or experiencing higher rates of intrusion, leading them to resort to overt aggression more frequently to defend their territory.

Plumage colouration does play a key role in mediating aggressive interactions between neighbors and floaters in a number of other territorial bird species, just as we have found in this study. In other avian species, individuals are less likely to approach or challenge more ornamented individuals or models (Pryke et al., 2001) and are more likely to challenge individuals with reduced or missing ornaments (Chaine & Lyon, 2008; Cline et al., 2016; Pryke & Andersson, 2003a, 2003b). Indeed, more ornamented territory holders experience lower rates of intrusion by conspecific males (Pryke & Andersson, 2003b; Pryke et al., 2001) while males with reduced ornaments experience much greater rates (Chaine & Lyon, 2008; Cline et al., 2016). Territorial males could receive intrusions from more distant neighbors who are prospecting for extra-pair mating opportunities or a higher quality territory. These individuals will be less familiar with each other and a signal of fighting ability could be beneficial in this context. While non-territorial “floaters” occur in island populations of song sparrows (Arcese, 1987), the number of floaters in our population is currently unknown and assessing the occurrence of floaters as well as the size and reflectance of their chest spotting would be helpful, since frequent interactions with floaters would make a signal associated with resource holding potential or aggression more likely to persist in a territorial species.

The findings in the current study support the hypothesis that spotting area is a signal used in male competitive interactions and influences receiver territorial aggression in song sparrows, although the mechanism of the association between the signal and aggression is unknown. To address this question, we are currently completing an experimental manipulation of the spotting area in male song sparrows and assessing the behavioural and hormonal consequences of these manipulations.

### 3-D printed models as a tool for behavioural assays

A second aim of our study was to assess the use of 3-D printed models as a replacement for taxidermic mounts in behavioural assays of aggression. While the use of 3-D printed models is becoming more common, relatively few studies have compared traditional taxidermic models and 3-D printed ones to ensure the 3-D model provides a biologically meaningful stimulus, particularly for behavioural studies (but see Bentz et al., 2019; Igic et al., 2015; Watson & Francis, 2015). We found our subjects in the urban areas responded to the taxidermic mount with greater territorial aggression than the 3-D printed models although they still responded to, and in some cases attacked, the 3-D printed models. The attack rates of birds which were tested with both 3-D models and the taxidermic mount were comparable (2 out of 11 birds attacked the mount, 3 out of 11 attacked the small-spotting 3-D model, and 1 out of 11 attacked the large-spotting 3-D model). Because we only had a single taxidermic mount, we cannot draw strong conclusions about the equivalence or lack thereof, of 3-D models and taxidermic mounts. Our results suggest researchers should validate their use of 3-D printed models by comparison to more realistic taxidermic mounts.

Though their validity across study systems still requires testing, 3-D printed models provide a very promising avenue for ecological research, including behavioural ecology. For intrusion experiments, the presence of a model presents a more realistic stimulus than song playback alone and can lead to stronger, and likely more realistic, behavioural and hormonal responses in some species (Chantrey & Workman, 1984; Wingfield & Wada, 1989). One major advantage of using 3-D printed models is the ability to manipulate shape, colour, posture, etc. of the visual stimuli and therefore gain more experimental control over treatments. Another advantage is that it is easy to 3-D print many models to reduce or completely avoid pseudoreplication without impacting wild populations by collecting specimens for taxidermic mounts (pseudoreplication was an issue in our study given we only had access to a single taxidermic mount). Indeed, 3-D printing has been used with great efficacy in several recent studies in birds (Fan et al., 2018; Igic et al., 2015; O’Connor, Brigham, & McKechnie, 2018) and other taxa (Watson & Francis, 2015), which suggests this technique can enhance a variety of ecological studies (Behm, Waite, Hsieh, & Helmus, 2018). We therefore believe that going forward 3-D printing will be a major benefit for behavioural ecology.

